# Entropic overcompensation of the N501Y mutation on SARS-CoV-2 S binding to ACE2

**DOI:** 10.1101/2022.08.30.505841

**Authors:** Natasha Gupta Vergara, Meghan Gatchel, Cameron F. Abrams

**Affiliations:** Department of Chemical and Biological Engineering, Drexel University, Philadelphia, Pennsylvania 19104, United States; Department of Biomedical Engineering, University of Delaware, Newark, Delaware, 19716, United States

## Abstract

Recent experimental work has shown that the N501Y mutation in the SARS-CoV-2 S glycoprotein’s receptor binding domain (RBD) increases binding affinity to the angiotensin-converting enzyme 2 (ACE2), primarily by overcompensating for a less favorable enthalpy of binding by a greatly reducing the entropic penalty for complex formation, but the basis for this entropic overcompensation is not clear [Prévost et al., *J. Biol. Chem*. (2021) 297;101151]. We use all-atom molecular dynamics simulations and free-energy calculations to qualitatively assess the impact of the N501Y mutation on enthalpy and entropy of binding of RBD to ACE2. Our calculations correctly predict that N501Y causes a less favorable enthalpy of binding to ACE2 relative to the original strain. Further, we show that this is overcompensated for by a more entropically favorable increase in large-scale quaternary flexibility and intra-protein root-mean squared fluctuations of residue positions upon binding in both RBD and ACE2. The enhanced quaternary flexibility stems from N501Y’s ability to remodel the interresidue interactions between the two proteins away from interactions central to the epitope and toward more peripheral interactions. These findings suggest that an important factor in determining protein-protein binding affinity is the degree to which fluctuations are distributed throughout the complex, and that residue mutations that may seem to result in weaker interactions than their wild-type counterparts may yet result increased binding affinity thanks to their ability to suppress unfavorable entropy changes upon binding.

## Introduction

Severe Acute Respiratory Syndrome Coronavirus 2 (SARS-CoV-2) is a novel virus that leads to the infectious disease Coronavirus 2019 (COVID-19). Initial transmission of SARS-CoV-2 involves binding of the S glycoprotein to sACE2 receptors on target cells. ^1^ SARS-CoV-2 S is a class-I trimeric viral fusion protein with each protomer comprising an S1 and S2 subunit (Fig. 1).^2^ S2 anchors S into the viral membrane and S1 is exposed on the membrane surface. The receptor binding domain (RBD) of S1 sub-unit binds to the ACE2 receptor on host cells. The ACE2 recognition epitope of RBD is hidden in the native timeric conformation, which much transition into an “RBD-up” conformation to allow sRBD:sACE2 complexation. This initial binding is the first step in membrane fusion and the energetically downhill cascade of events that lead to host cell entry.^3^ RBD is the locus of several point mutations that have led to the rise of many new SARS-CoV-2 variants.

**Figure 1:**
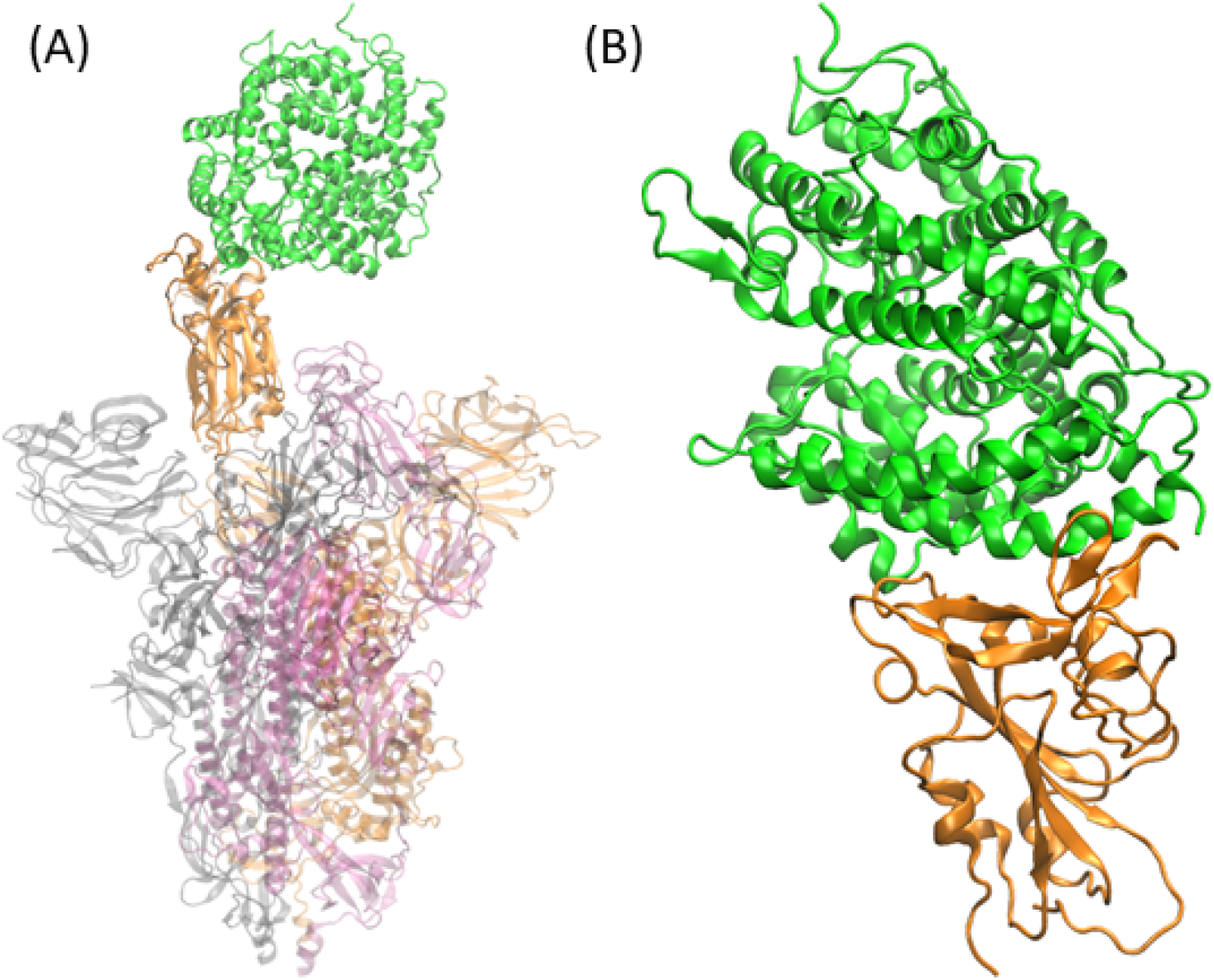
(A) SARS-CoV-2 S Glycoprotein S trimer with ACE2 (green) bound to one sRBD (orange) in the up position. (B) sACE2 bound to sRBD as modeled in this work (from PDB 6m0j^4^).

One such point mutation is the change from asparagine to tyrosine at position 501 (N501Y). X-ray crystallographic structures shows that 501 interacts directly with residues on ACE2.^4^ N501Y is present in many variants, including Alpha (B.1.1.7), Beta (B.1.351), Gamma (P.1), and Omicron (B.1.1.529).^5,6^ The N501Y mutation has been shown to increase sACE2-sRBD binding affinity^7,8^ as well as infectiousness and transmissibility of SARS-CoV-2.^9^ Using isothermal titration calorimetry on soluble versions of RBD and ACE2, Prévost et al. showed that N501Y_sRBD_ leads to *less* favorable enthalpy of binding which is more than offset by *more* favorable entropy of binding, which both increases the overall sRBD:sACE2 binding affinity and suppresses the decrease in binding affinity with an increase in temperature, all relative to the Wuhan strain. The basis for this “entropic overcompensation” is not clear. The competing roles of enthalpy and entropy changes upon binding to binding affinity are well-appreciated, but attribution of such remarkably large changes to both to one residue in a protein protein interface can be challenging.^10^ To attempt an explanation, we have conducted a series of all-atom molecular dynamics (MD) simulations of the soluble form of RBD (sRBD), the soluble form of ACE2 (sACE2), and their complexes. We evaluate the enthalpy of binding for both the Wuhan and N501Y strains, as well as the impact of binding on the conformational flexibility of the complex and of each protein. Our results allow us to postulate a structural rationale for the N501Y mutation’s effect on both the enthalpy and entropy of binding consistent with experimental results.

## Methods

For this work we modeled five distinct systems: sRBD, sRBD_N501Y_, sACE2, sRBD:sACE2, and sRBD_N501Y_:sACE2. For purposes of distigushing between systems, we will refer to those systems in which the sRBD sequence is that of the Wuhan strain as “wild-type” and those with the N501Y_sRBD_ mutation as “mutant”. All atomic coordinates were based on the PDB entry 6M0J.^4^ All systems were solvated in explicit TIP3P water and neutralized with counterions. Equilibrium MD simulations were run using NAMD for 40 ns. ^11^ Three replicas of each system were simulated. All systems utilized periodic boundary conditions, a nonbonded cutoff of 11 Å, a particle-mesh Ewald grid spacing of 1 Å, and a timestep of 2 fs. All simulations were conducted at 310 K using a Langevin themostat with a friction parameter of 5 ps^*−*1^. Molecular-Mechanics/Generalized Born Surface Area (MM/GBSA) free-energy calculations were conducted using VMD ^12^ to evaluate the enthalpic contribution to binding free energy.

Entropic contributions were estimated using carma. ^13^ Heavy atoms in residues at the sRBD:sACE2 interface were selected to fit corresponding trajectories in both the WT and N501Y_sRBD_ mutation and an approximation of the upper limit to the absolute entropy was calculated using Schlitter’s formula.^14^

Total hydrogen bond occupancy and interatomic distances were evaluated using VMD. ^12^ To identify hydrogen bonds, a distance-based method was used where we identify two residues H-bonded when the distance between their heavy atoms is ≤3.3 Å over 30 ns of equilibrated MD trajectory per replica. All heavy atoms at the interface of each system was assessed with the respective nearest neighbor distance atomic distance for the partner residue. Normal mode analysis was conducted using Bio3d^15^ and visualized in Pymol. ^16^

## Results

### Binding thermodynamics

We first demonstrate that our simulations provide a qualitatively correct comparison of the thermodynamics of sRBD:sACE2 binding between WT and N501Y systems. In Table 1, we report the enthalpic contribution computed using MM/GBSA, denoted Δ*H*_MM*/*GBSA_, and the entropic contribution to the free energy of binding calculated via Schlitter’s formula over sRBD:sACE2 interface heavy atoms, denoted −*T* Δ*S*^s^, both in kcal/mol, for both the WT and mutant sRBDs. Consistent with the experimental findings of Prévost et al., ^7^ reported for reference in Table 1, we see that the effect of the N501Y_sRBD_ mutation is to make the enthalpy of binding less favorable while making the entropy of binding more favorable. We stress that we are not claiming that the Schlitter-based entropy change over interface residues constitutes the total entropic contribution to binding; only that it suggests that entropic contributions from the residues at the interface are consistent with the experimental observation that −*T* Δ*S* is lower for N501Y_sRBD_ than for WT. The Schlitter entropy includes contact residues from both sRBD and sACE2. With this correspondence between simulations and experiments establish, we next explain the bases for these thermodynamic differences.

**Table 1:** MM/GBSA enthalpy of binding (Δ*H*_MM*/*GBSA_) and Schlitter’s entropy of binding among interface residues (−*T* Δ*S*^s^) WT and N501Y_sRBD_ systems at 310 K, computed from MD simulations. For reference, enthalpies and entropies from isothermal titration calorimetry experiments at 310 K of Prévost et al. ^7^ are shown (Δ*H* and −*T* Δ*S*, respectively).

| System | This work (simulations) |  | ITC<sup>7</sup> |  |
| --- | --- | --- | --- | --- |
| | $\Delta H_{\text{MM/GBSA}}$ (kcal/mol) | $-T\Delta S^s$ (kcal/mol) | $\Delta H$ (kcal/mol) | $-T\Delta S$ (kcal/mol) |
| WT | $-30.22 \pm 3.54$ | $22.03 \pm 7.39$ | -22.9 | 12.4 |
| N501Y | $-27.63 \pm 1.29$ | $18.11 \pm 5.49$ | -21.6 | 10.0 |

### Enthalpic effects of the N501Y mutation

We characterized enthalpic contributions to binding by enumerating and classifying residue-residue interactions at the protein-protein interface as depicted in Fig. 2. For all residue pairs at the interface, distributions of nearest interatomic distance in each pair were calculated. A instantaneous residue-residue contact is defined as any pair of residues for which at least one atom on the first is within 6 Å of at least one atom on the second. Although we focus on the interface, we tabulated interactions both across the RBD:ACE2 interface and within either RBD or ACE2. Fig. 3 depicts the most significant interresidue interactions at the interface, defined as those residue pairs which showed the largest change in nearest interatomic distance between the WT and mutant systems. This figure also serves to illustrate the extended nature of the interface which prompts us to classify the interactions between “peripheral” and “central”. We now turn to detailed consideration of each set of residue pairs.

**Figure 2:**
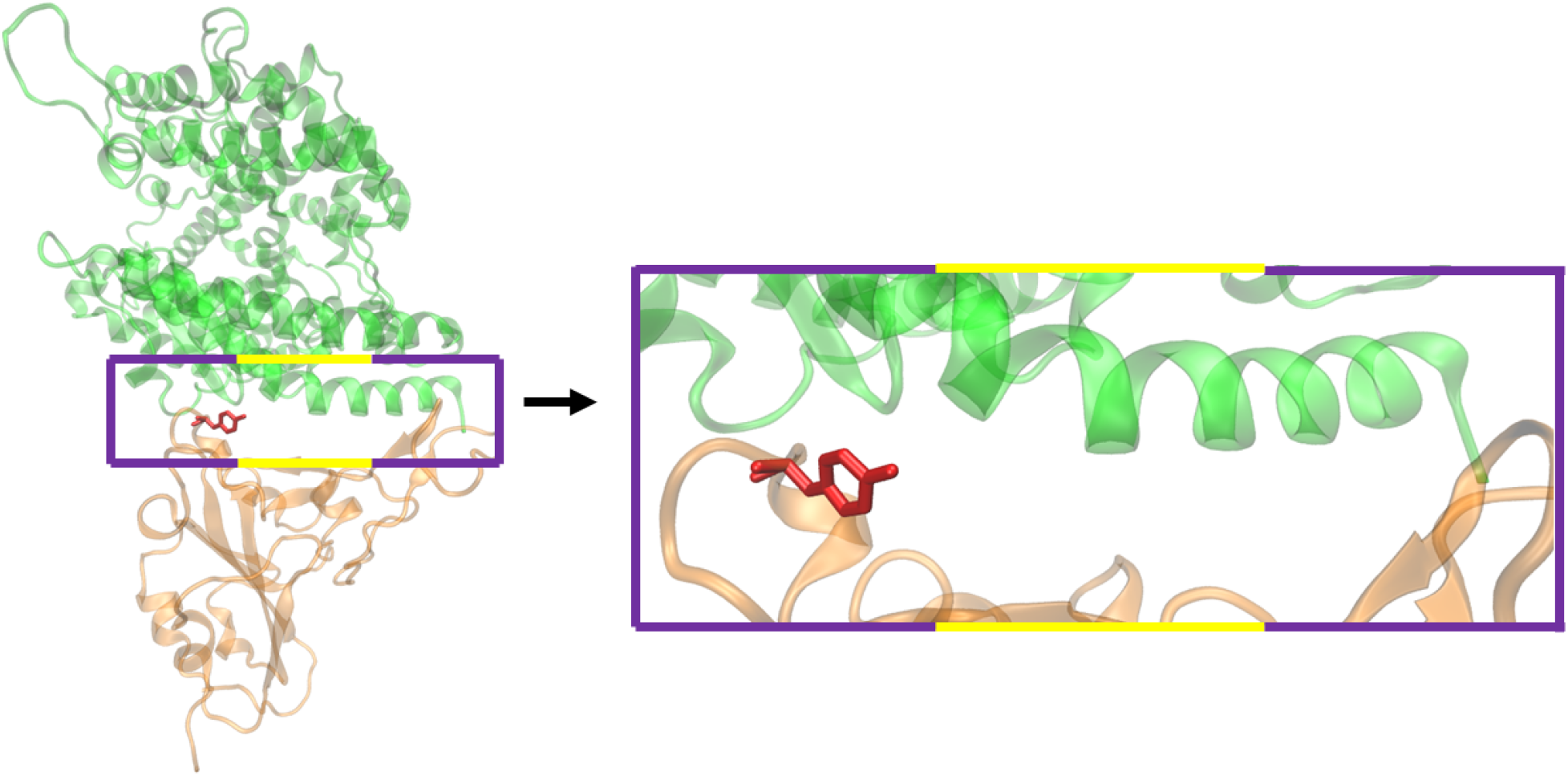
Section of the sRBD:sACE2 interface that is evaluated for residue contacts. N501 is shown in red. Peripheral interactions are outlined in purple while central interactions are outlined in yellow.

**Figure 3:**
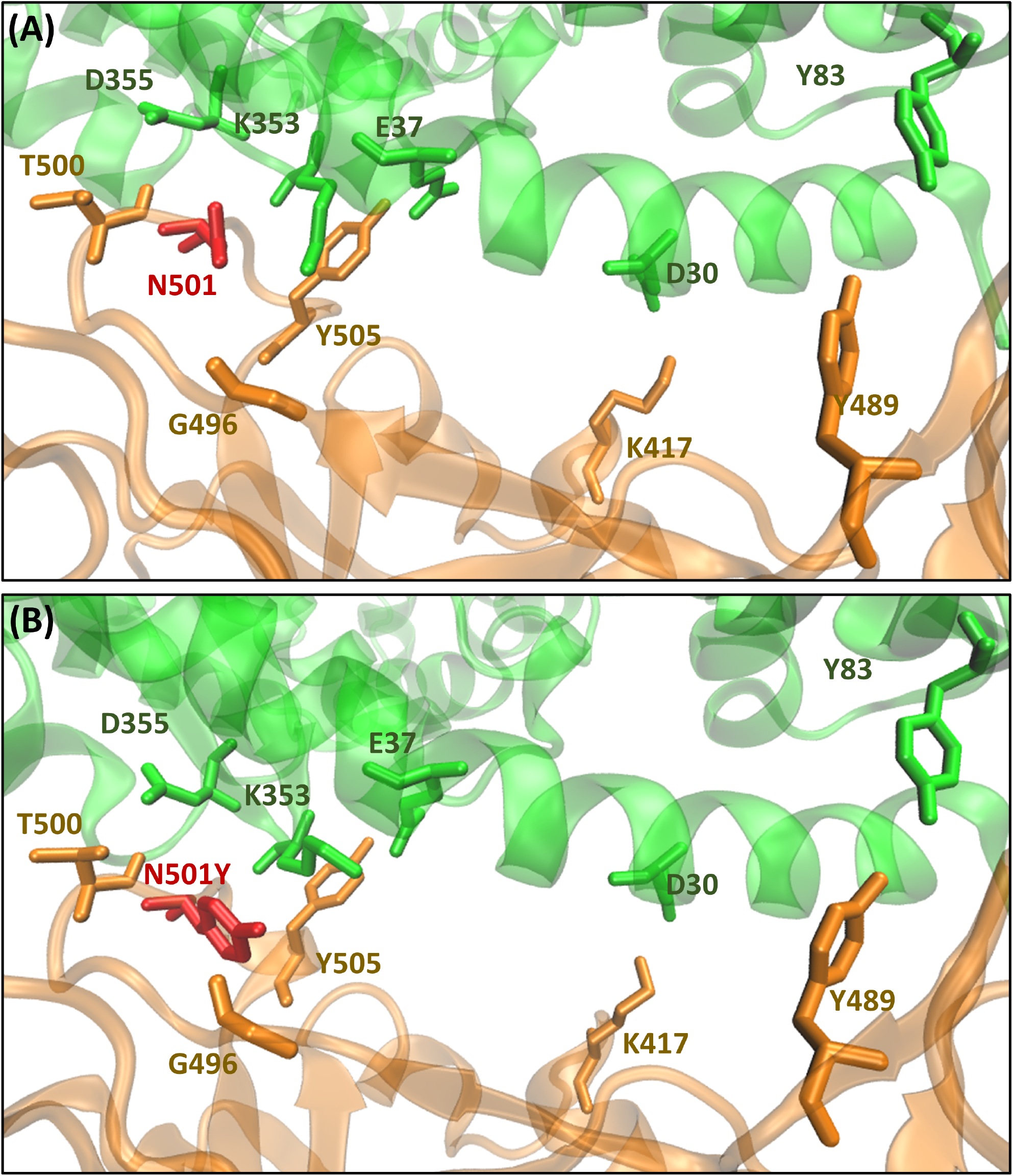
Detailed views of the sRBD:sACE2 interface in representative configurations from the MD simulations; (A) WT, (B) N501Y. In both panels, residue 501_sRBD_ is shown in red licorice, and sRBD in red cartoon and sACE2 in green cartoon.

The first effect of N501Y on the interface structure involves the ACE2 residue K353. Fig. 4 shows the interactions that K353_sACE2_ makes with residues in sRBD in both the WT and mutant systems. Fig. 4(A) and (C) show how K353_sACE2_ switches from H-bonding to the backbone carbonyl of G496_sRBD_ in WT to H-bonding to the side-chain hydroxyl of Y501_sRBD_ in the mutant. Both of these H-bonds are fairly well occupied, as evidenced by the interatomic density distributions compiled from the final 30 ns of all MD simulations, shown in Fig. 4(C) and (D). Note that this switch of interaction partners of K353_sACE2_ moves an RDB:ACE2 contact from a more central location to a more peripheral location on the interface.

**Figure 4:**
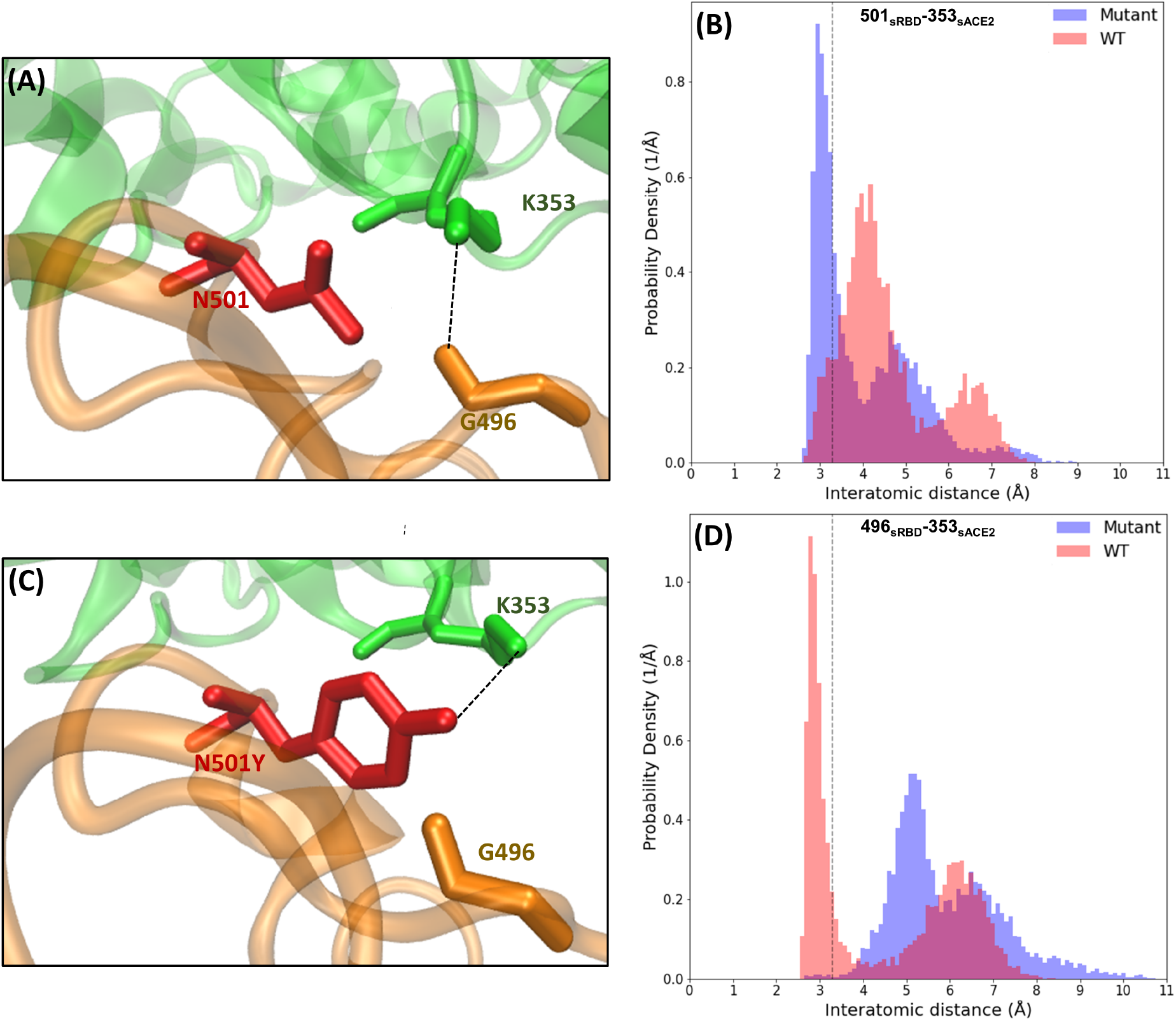
Close up views of interactions among 501_sRBD_, G496_sRBD_, and K353_sACE2_ for WT (A) and mutant (C); sRBD is shown in orange and sACE2 in green, and dashed lines indicate H-bonds. (B) Probability density of closest interatomic distances between residues 501_sRBD_ and 353_sACE2_ from MD simulations of the WT (red) and mutant (blue) systems. (D) Probability density of closest interatomic distances between residues 496_sRBD_ and K353_sACE2_ from MD simulations of the WT (red) and mutant (blue) systems.

The mutant also displays other more peripheral interactions. Fig. 5 shows that the H-bond accepted by D355_sACE2_ from the backbone amide of T500_sRDB_ is significantly more populated in the mutant than in WT. Fig. 6 shows that the formation of the K353_sACE2_-Y501_sRBD_ H-bond in the mutant sacrifices a more central H-bond between E37_sACE2_ and Y505_sRBD_. The mutant also sacrifices a small amount of occupancy of a central H-bond between D30_sACE2_ and K417_sRBD_. Finally, in Fig. 7, we show that the mutant acquires an intermittent peripheral H-bond between the hydroxyls on Y83_sACE2_ and Y489_sRBD_. The general pattern of changes in H-bonding interactions between sACE2 and sRBD upon the N501Y_sRBD_ mutation seems to be reduction in more centrally located H-bonds to more peripherially located H-bonds. We will see in the following sections some large-scale conformational analysis that may provide more insight into what implications this observation may have on the motion of the complex at equilibrium. All in all, the overall occupancy of H-bonds is slightly higher in WT than in the mutant (9%), consistent with the observation that the enthalpy of binding is somewhat less favorable for the mutant system.

**Figure 5:**
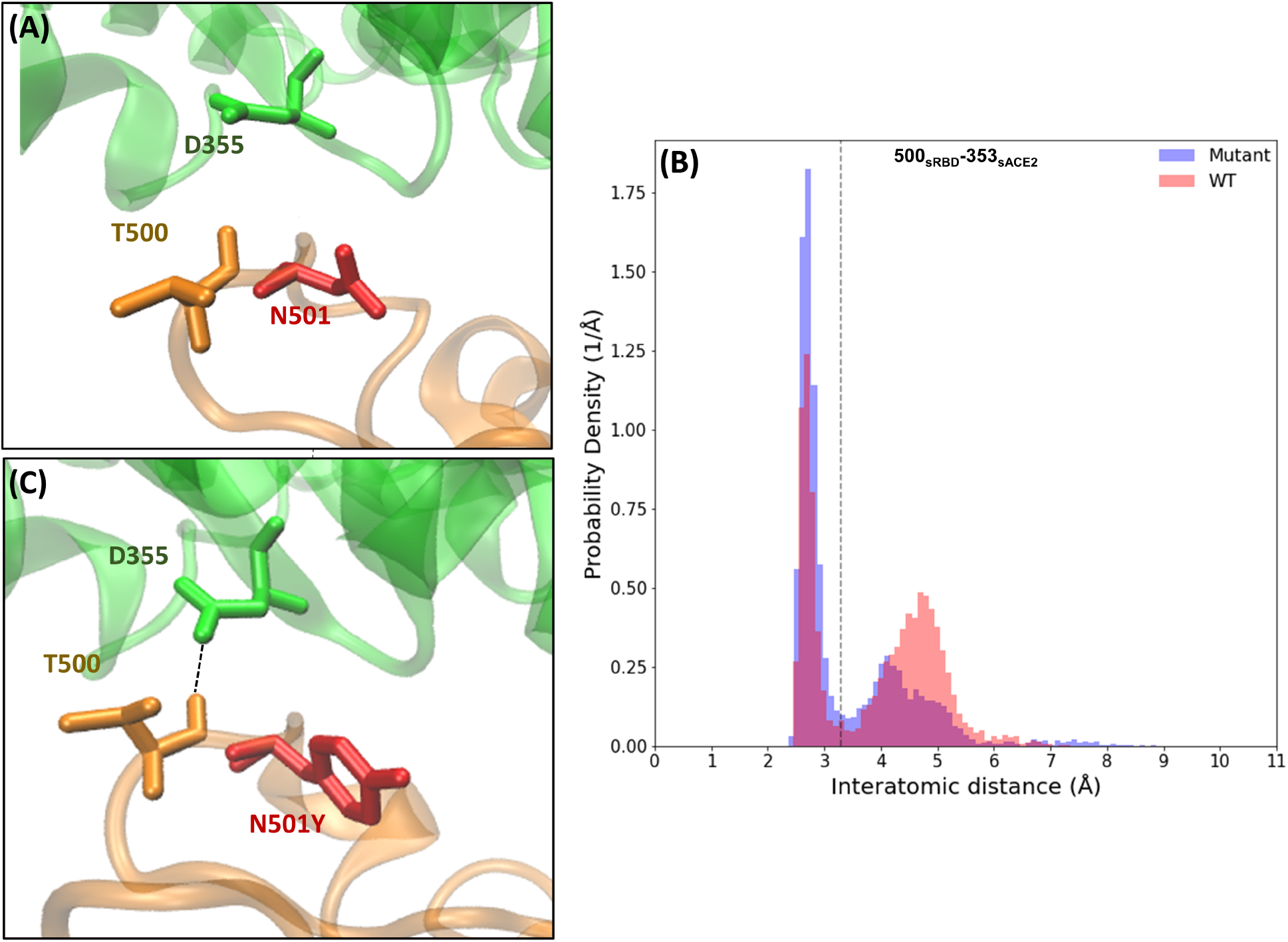
Close up views of interactions among 501_sRBD_, T500_sRBD_, and D355_sACE2_ for WT (A) and mutant (C); sRBD is shown in orange and sACE2 in green, and dashed lines indicate H-bonds. (B) Probability density of closest interatomic distances between residues T500_sRBD_ and D355_sACE2_ from MD simulations of the WT (red) and mutant (blue) systems.

**Figure 6:**
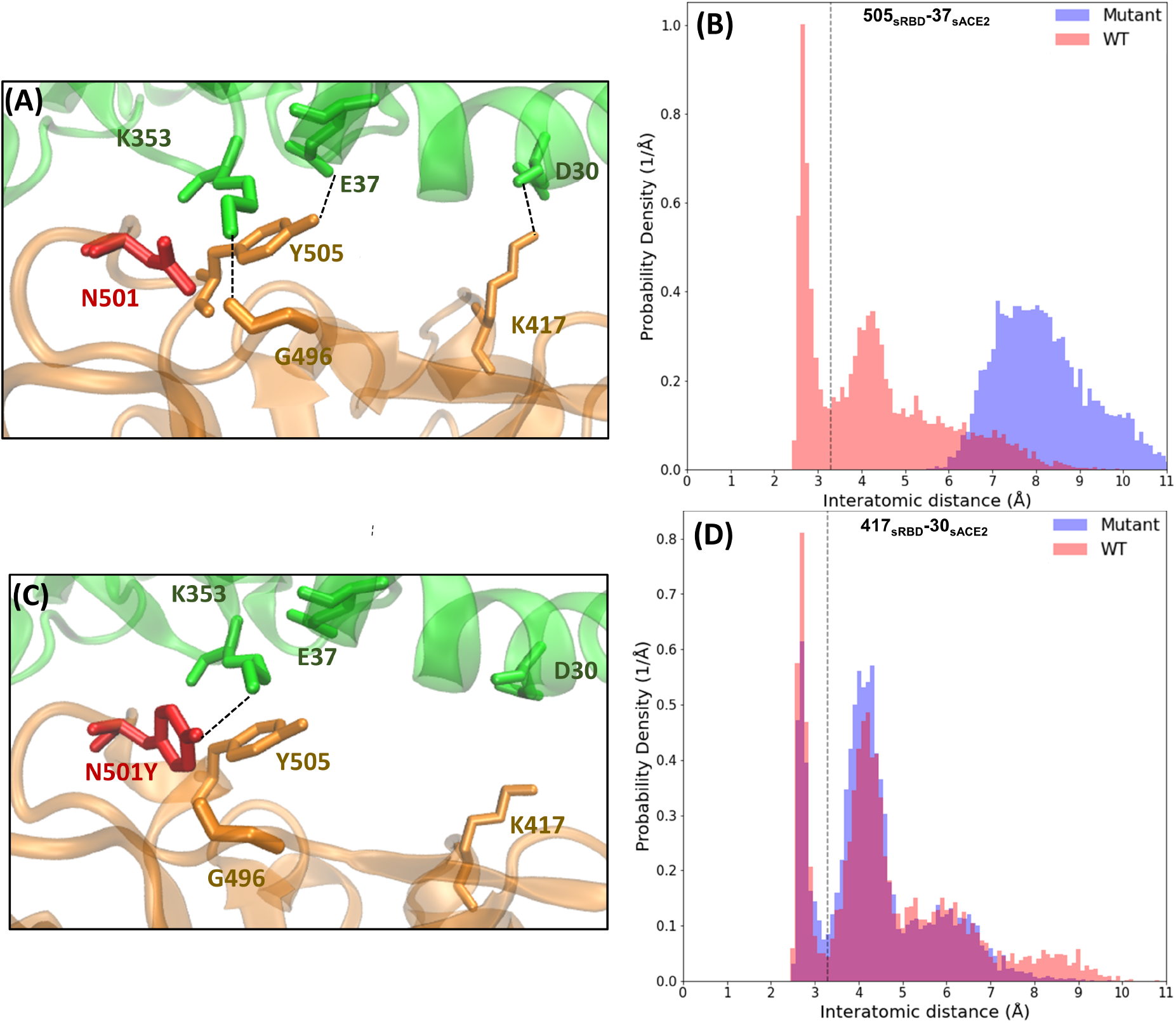
Close up views of interactions among Y505_sRBD_ and E37_sRBD_, and K417_sRBD_ and D30_sACE2_ for WT (A) and mutant (C); sRBD is shown in orange and sACE2 in green, and dashed lines indicate H-bonds. Interactions previously shown among 501_sRBD_, G496_sRBD_, and K353_sACE2_ are shown here again for reference. (B) Probability density of closest interatomic distances between residues Y505_sRBD_ and E37_sACE2_ from MD simulations of the WT (red) and mutant (blue) systems. (D) Probability density of closest interatomic distances between residues K417_sRBD_ and D30_sACE2_.

**Figure 7:**
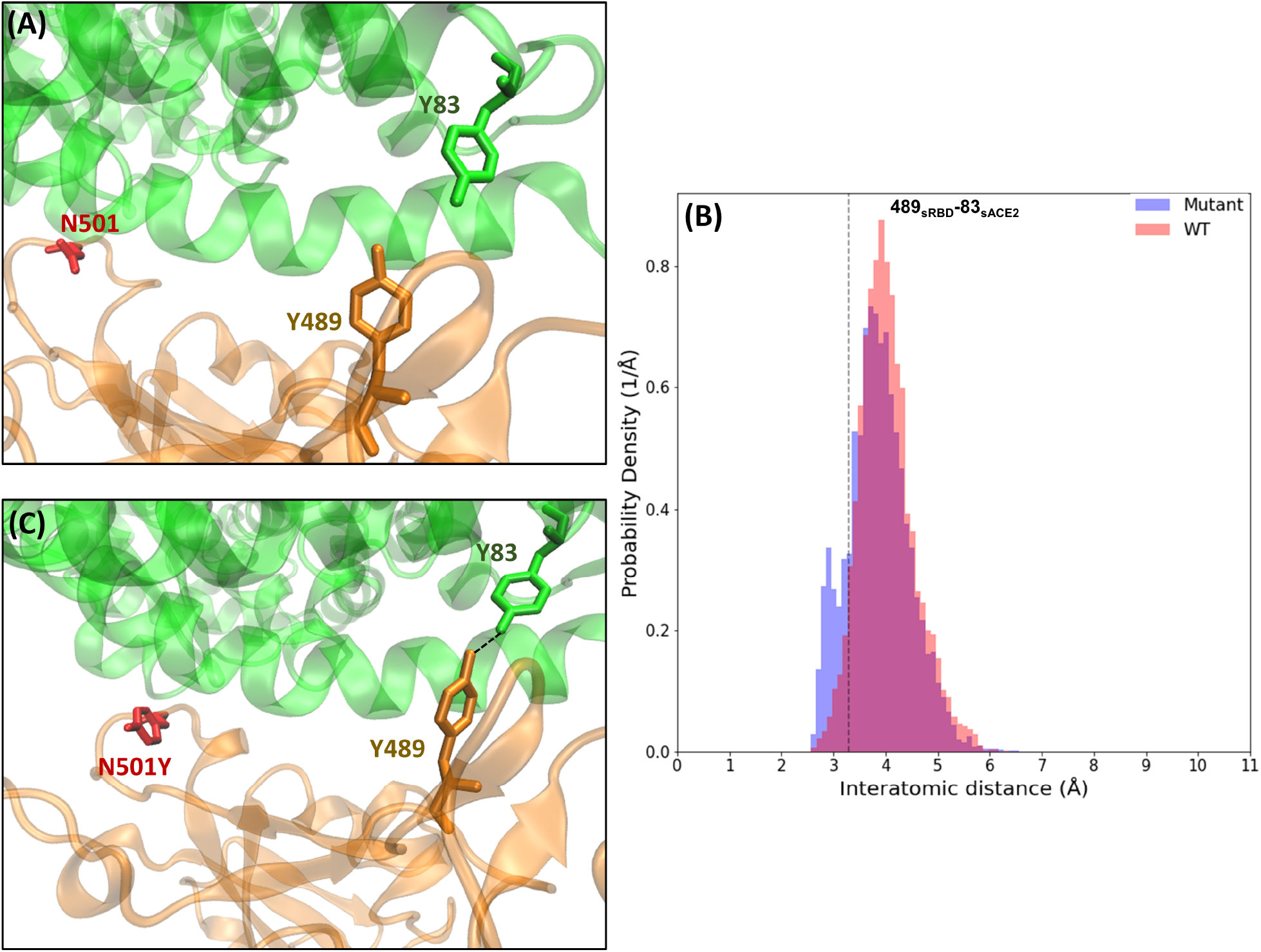
Close up views of interactions among Y489_sRBD_ and Y83_sRBD_ for WT (A) and mutant (C); sRBD is shown in orange and sACE2 in green, and dashed lines indicate H-bonds, and 501_sRBD_ is highlighted for reference. (B) Probability density of closest interatomic distances between residues Y489_sRBD_ and Y83_sACE2_ from MD simulations of the WT (red) and mutant (blue) systems.

### Entropic effects of the N501Y mutation

To understand how the N501Y mutation might entropically affect binding, we first investigated interatomic positional fluctuation correlations. Dynamic cross-correlation matrices were computed from all aggregate replica trajectories for the WT and mutant systems. Each matrix element *c*_*ij*_ for atoms *i* and *j* is computed via

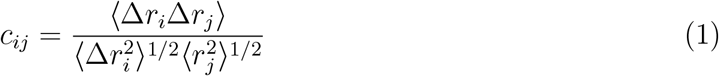

where Δ*r*_*i*_ = *r*_*i*_ − ⟨*r*_*i*_⟩ is the instantaneous fluctuation in cartesian coordinates of atom *i*. Graphical representations of the WT and mutant cross-correlation matrices appear in Fig. 8. Residues corresponding to RBD and ACE2 are denoted along each axes. The positive values of *c*_*ij*_ are shown in red and the negative values are shown in blue. From this we observe that there are fewer overall correlations between sRBD and sACE2 in the mutant system than in WT. ACE2 to ACE2 correlation is also less pronounced in the mutant than in WT. From this we infer that the change in the layout of interprotein H-bonds upon mutation is associated with weaker dynamic coupling between the two proteins, while at the same time sacrificing very little enthalpy of binding. A different view of the interatomic correlations is shown in Fig. 9, where we render line segments, colored by correlation value, between atoms for which |*c*_*ij*_| ≥ 0.4. These help to illustrate that the WT complex displays much more correlated fluctuations across the protein-protein interface than does the mutant.

**Figure 8:**
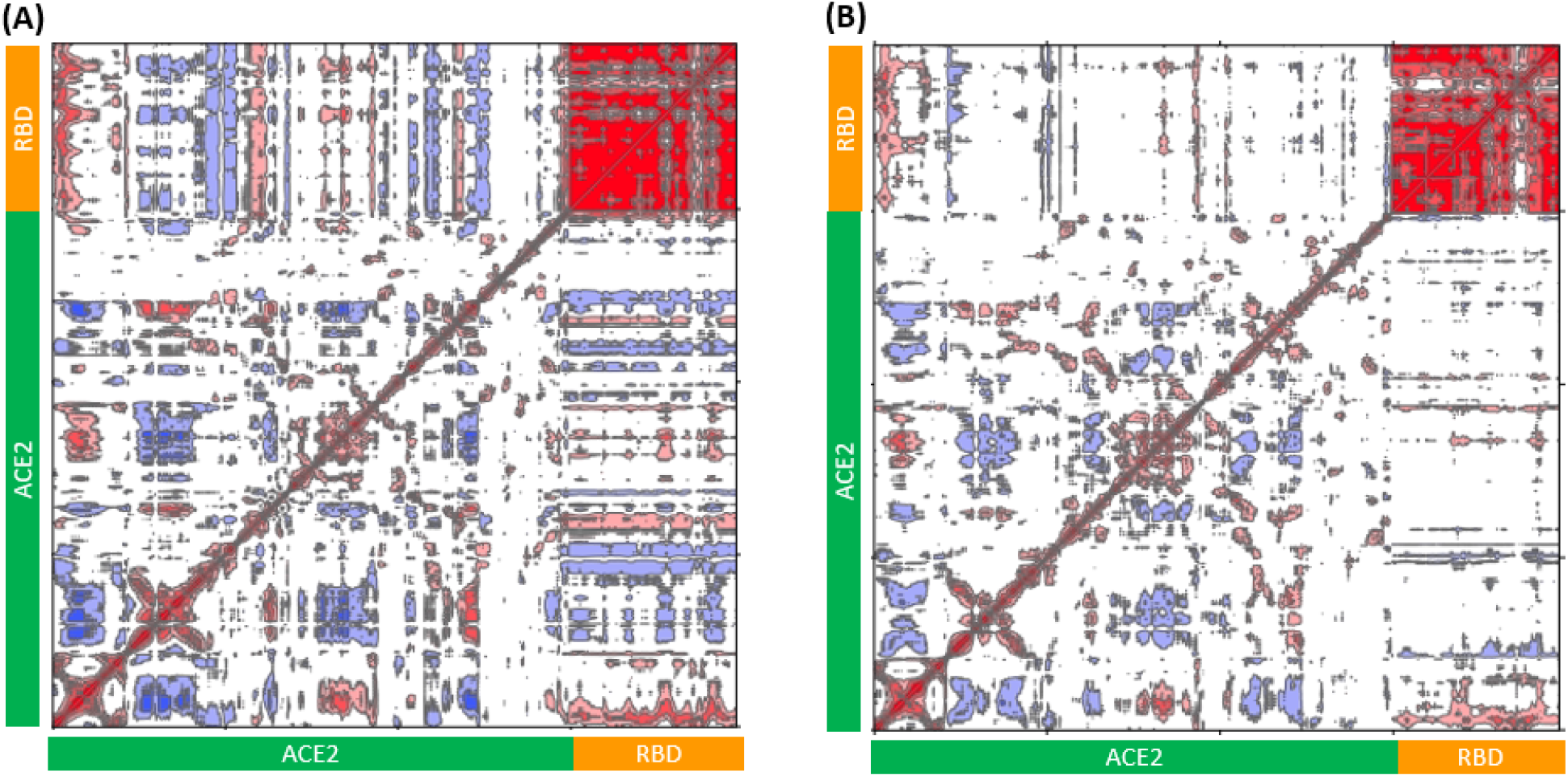
(A) Dynamic cross-correlation matrix of WT sRBD and sACE2. (B) Dynamic cross-correlation matrix of the N501Y_sRBD_ mutant sRBD and sACE2. Red represents correlated and blue represents anticorrelated motions.

**Figure 9:**
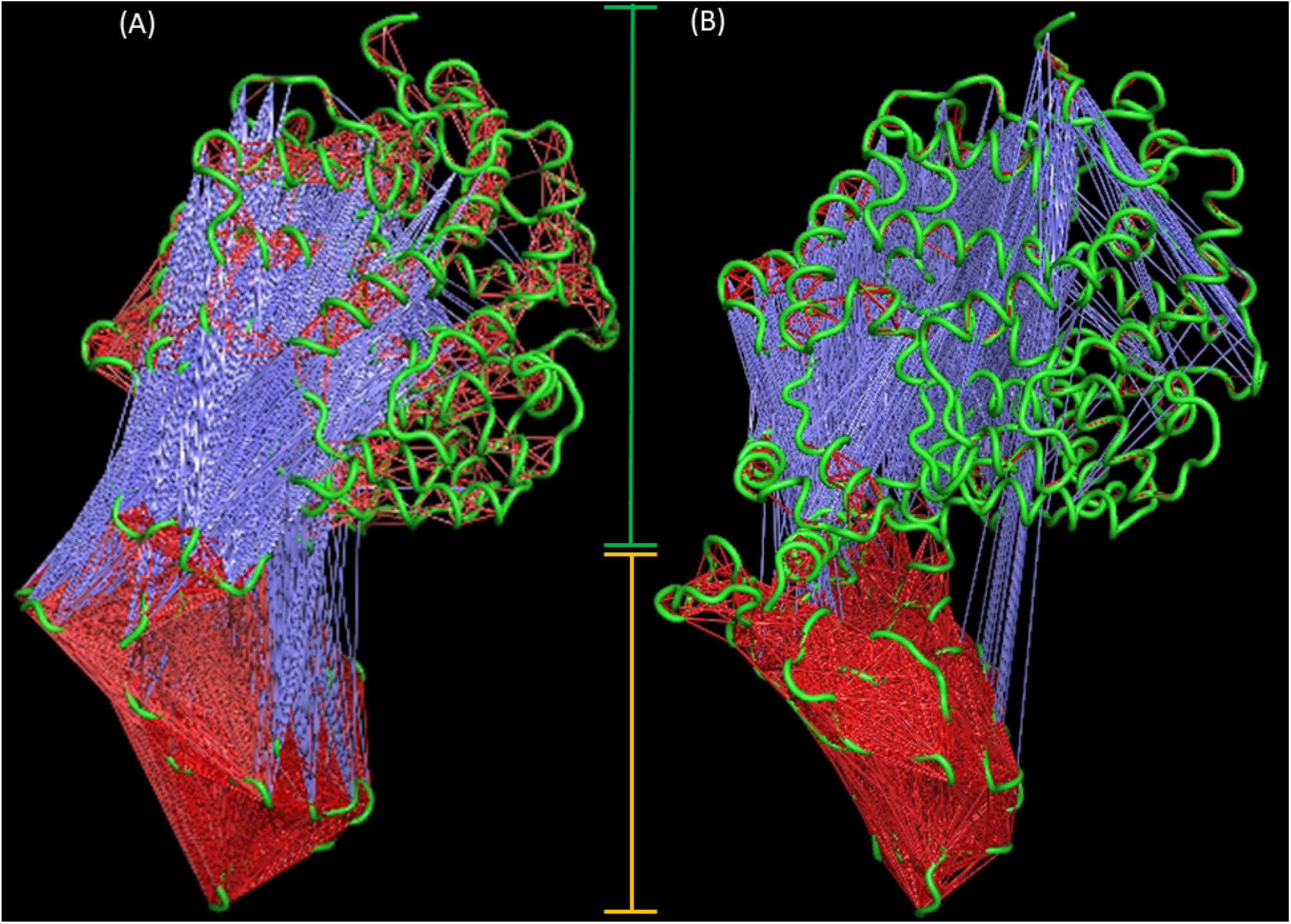
Mapping atomic correlations onto the sRBD:sACE2 complex, where atom pairs for which |*c*_*ij*_| ≥0.4 are indicated with lines with colors indicated magnitude as in Fig. 8. (A) WT; (B) Mutant. The subunit corresponding to ACE2 region identified by a green vertical line and RBD is identified by the orange vertical line.

Normal mode analysis (NMA) reveals that these changes in correlations reduce primarily to large scale relative motion of the two proteins, which is much more pronounced in the mutant than in WT. The lowest-frequency nontrivial normal mode is depicted in Fig. 10 in a vector-field representation. Each vector represents the direction of motion of the residues where the length indicates the vibrational amplitude. From this we see that the mutant complex displays much more large-scale flexibility, with a bending motion about the protein:protein interface. The supplementary movie showing the first non-trivial mode in each system shows that there is a hinge motion at the binding interface that shows increased motion for mutant in comparison to WT. Although qualitative, this interpretation also supports the observation that the entropy of binding of the mutant is more entropically favorable than WT.

**Figure 10:**
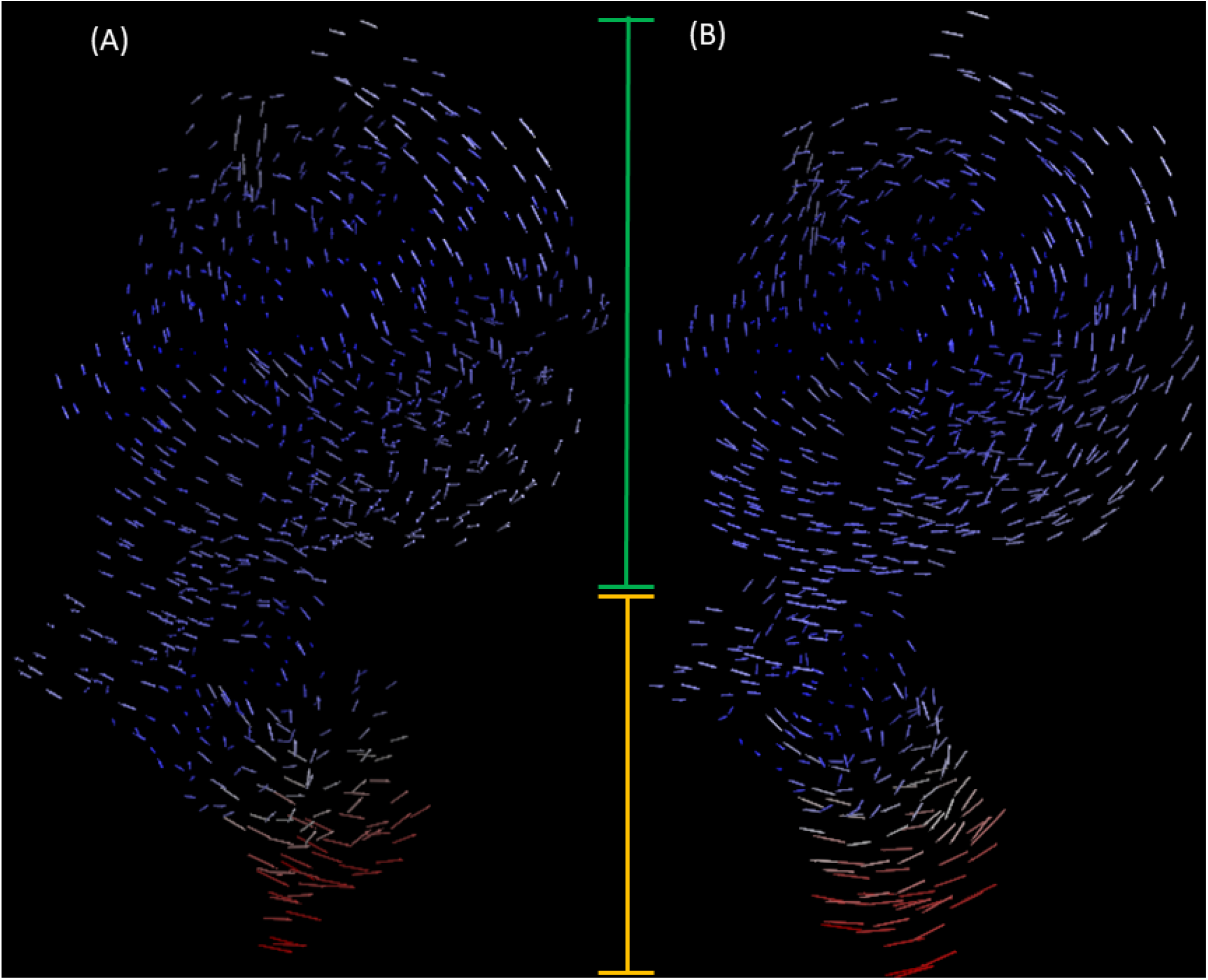
Vector field representation of the first nontrivial normal mode in (A) WT and (B) mutant. The subunit corresponding to ACE2 region identified by a green vertical line and RBD is identified by the orange vertical line.

NMA argues that one entropic effect of the mutation is the population of a large-scale flexibility mode in the complex that is not populated in the WT, and the H-bonding patterns seem to suggest that the more peripheral layout of H-bonds underlies this flexibility. However, the effect of the mutation is felt across the spectrum of normal modes. Because higher-frequency modes typically correspond to more localized fluctuations, we opt to consider root mean squared fluctuations (RMSF) of residue positions. Fig. 11 shows RMSF across the entire complex for both systems, and we observed that the mutant is more flexible overall across both ACE2 and RBD. In Fig. 12 we show the RMSFs of each protein from separate *apo* simulations together with the protein-specific RMSFs from the complex simulations, to isolate the effects of the mutation on the internal flexibility of each protein. These differences are revealed by taking the difference of the latter with the former, yielding ΔRMSFs shown in Fig. 12(C). Here we observe generally more negative ΔRMSF for the mutant than for WT, indicating slightly more suppression of fluctuations upon binding for the mutant. Residues with negative values of ΔRMSF seem to include, as expected, interface binding residues.

**Figure 11:**
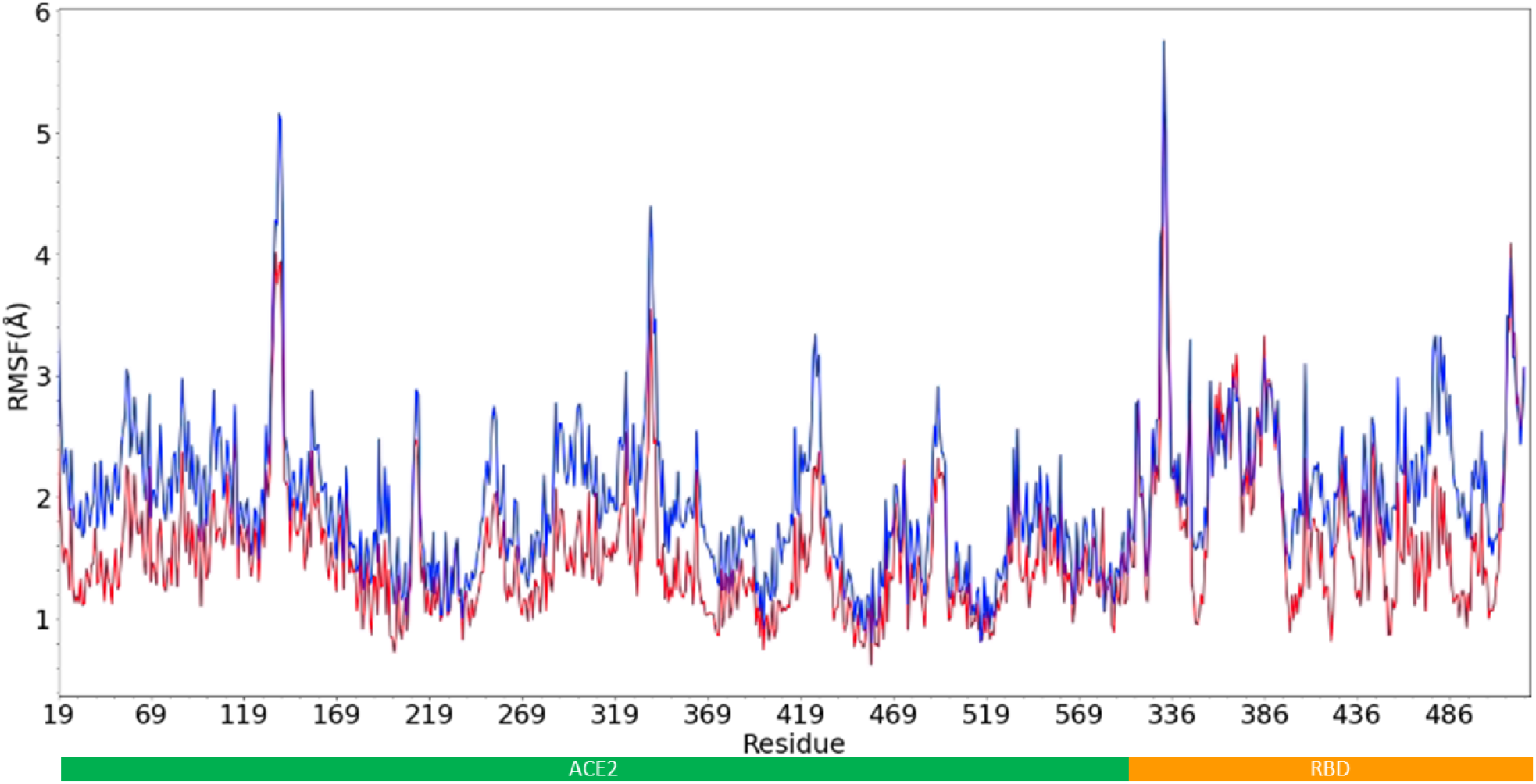
(A) RMSF of sACE2 and sRBD system for WT (red) and N501Y_sRBD_ mutant systems (blue).

**Figure 12:**
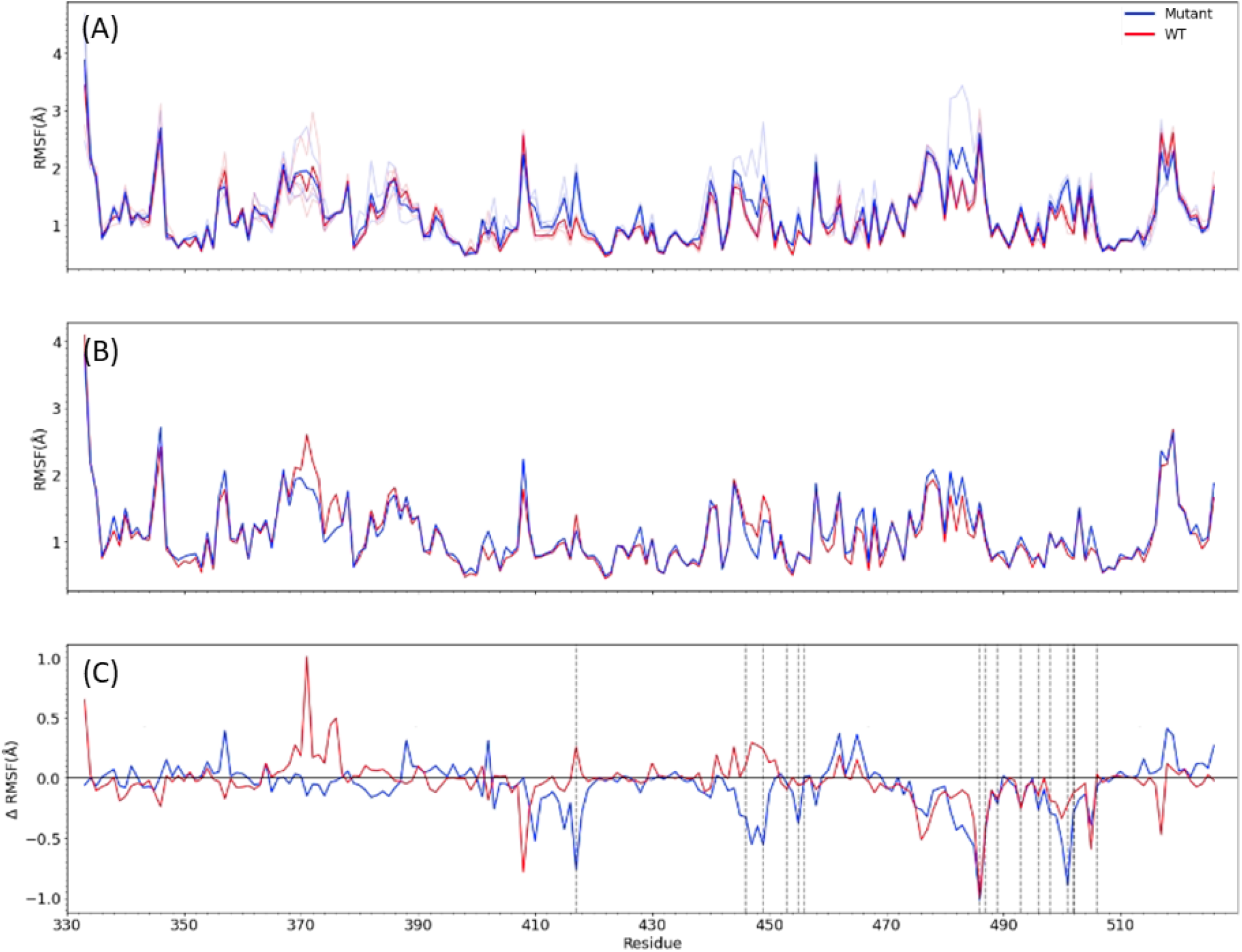
(A) RMSF of sRBD for apo WT (red) and N501Y_sRBD_ (blue) mutant systems. (B) RMSF of sRBD in complex with sACE2 (aligned on sRBD) for WT (red) and N501Y_sRBD_ (blue) systems. (C) ΔRMSF for sRBD, defined as RMSF of sRBD in complex with sACE2 minus RMSF of apo sRBD, for WT (red, mean: -0.03 Å) and N501Y_sRBD_ (blue, mean: -0.07 Å) systems. Residues that bind to sACE2 are indicated with dashed vertical lines.

The apparent greater degree of suppression of fluctuations upon binding for the mutant relative to WT seems counter to the observation that the magnitude of the entropy change of binding is *lower* for mutant. However, fluctuations in sRBD alone do not tell the whole story. In Fig. 13 we show impact of sRBD binding on the sACE2 RMSF. The ΔRMSF for sACE2 shows significantly more positive values for the mutant than for WT, indicating that fluctuations in sACE2 increase upon binding to mutant sRBD.

**Figure 13:**
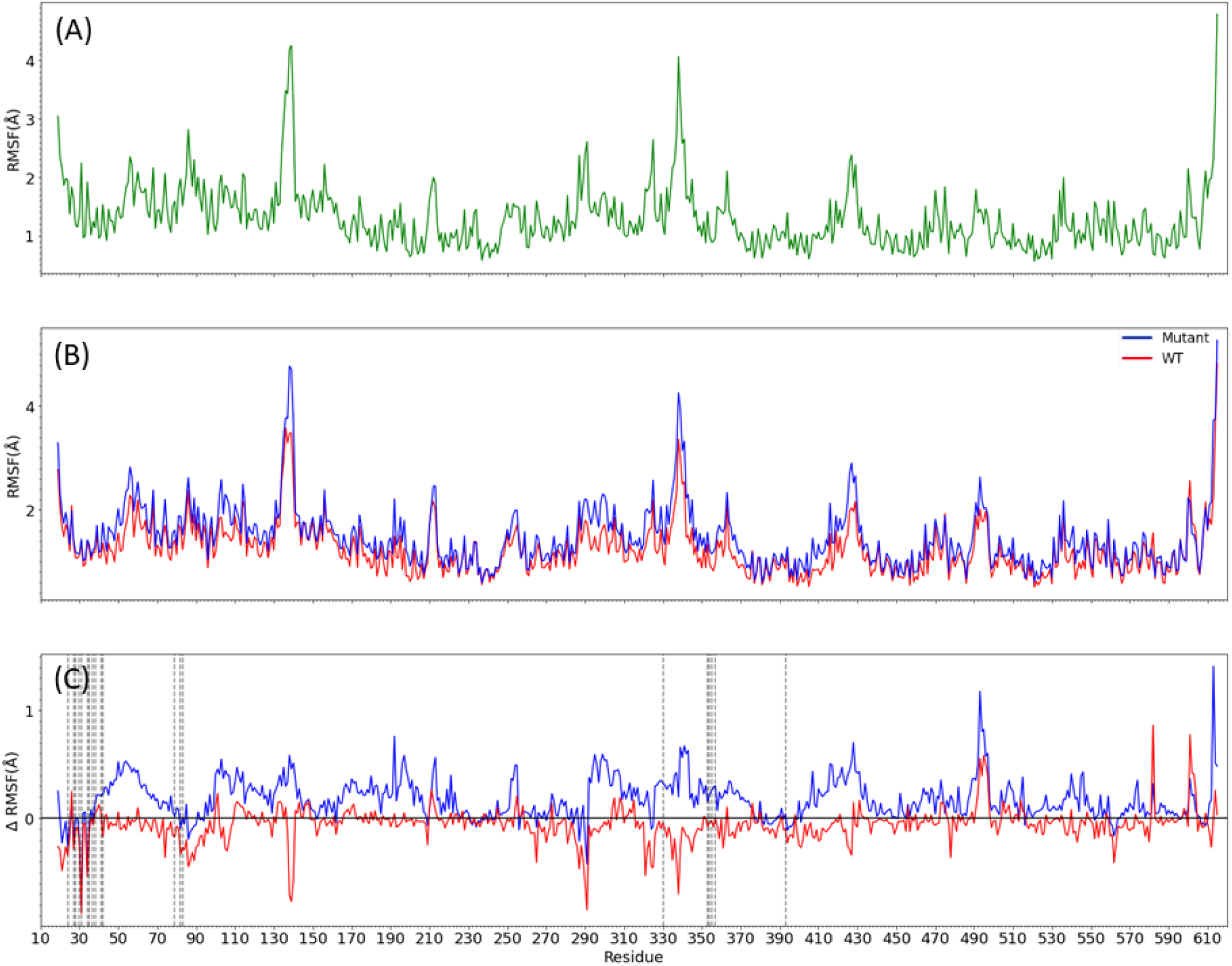
(A) RMSF of sACE2 from an apo simulation. (B) RMSF of sACE2 in complex with sRBD (aligned on sACE2) for WT (red) and N501Y_sRBD_ (blue) systems. (C) ΔRMSF for sACE2, defined as RMSF of sACE2 in complex with sRBD minus RMSF of apo sACE2, for WT (red, mean: -0.06 Å) and N501Y_sRBD_ (blue, mean: 0.18 Å) systems. Residues that bind to sRBD are indicated with dashed vertical lines.

The ΔRMSF was mapped onto the crystal structure of the sRBD in Fig. 14 to better visualize where the ΔRMSF values are located on the sRBD in relation to sACE2. We see that the majority of the more negative ΔRMSF values are at the interface of sACE2-sRBD for the mutant N501Y_sRBD_ system. Note that the rest of the sRBD has more positive ΔRMSF in the mutant system, indicating that although the contact residues may be more stable, the sRBD itself gains more flexibility upon binding. The overall picture is therefore one in which mutant sRBD fluctuations are seemingly transferred to sACE2 upon binding. Fig. 14 shows structures of the complexes colored by each protein’s respective ΔRMSF, and it is easy to see that in the mutant systems, although sRBD fluctuates *less* upon binding near the interface, sACE2 fluctuates more throughout the protein.

**Figure 14:**
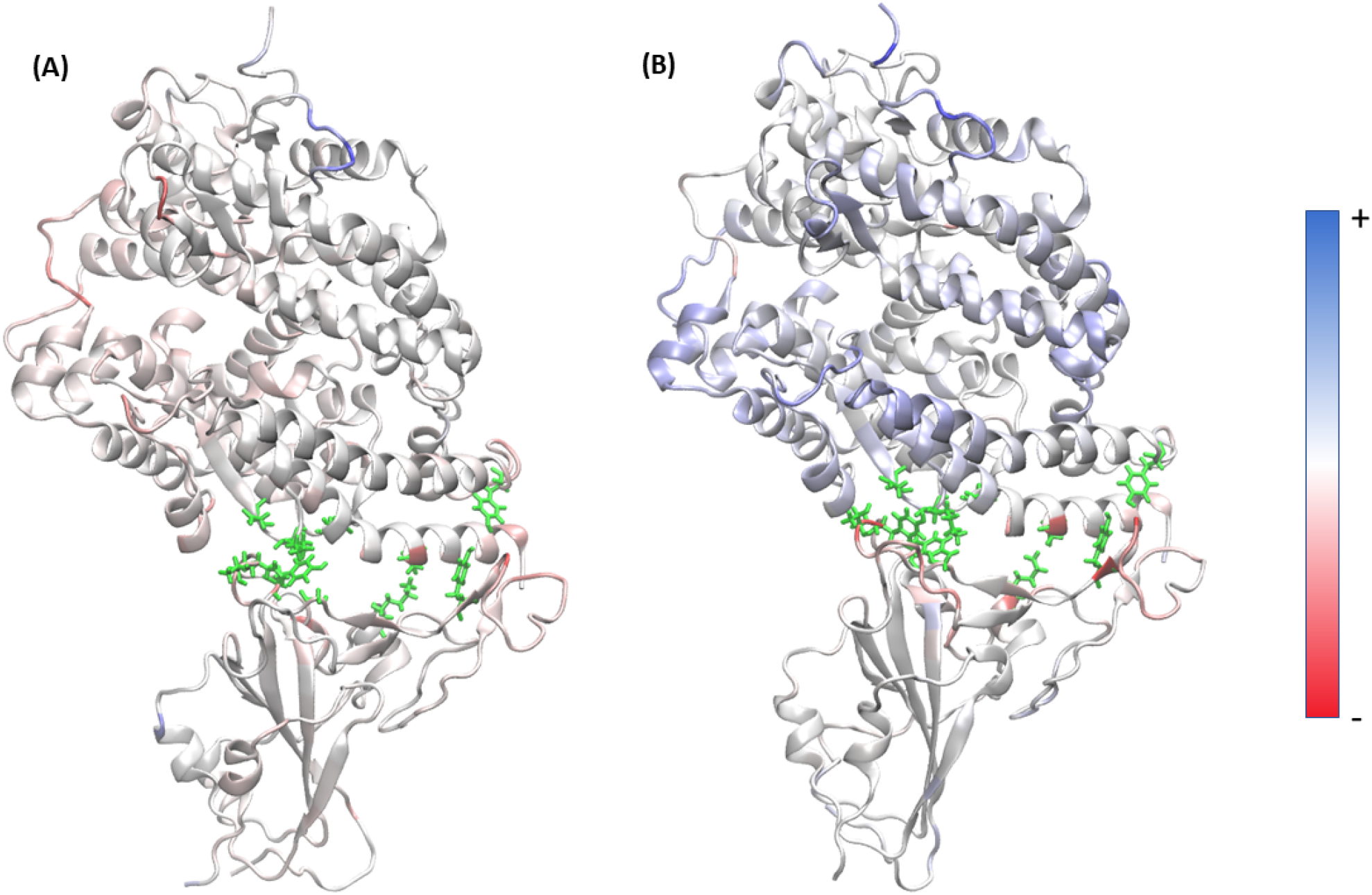
Cartoon representations of the sRBD:sACE2 complex, colored by ΔRMSFs, with redder colors more negative and bluer more positive, for (A) WT sRBD and (B) N501Y. Positive values correspond to increased flexibility upon binding while negative values correspond to a decrease in stability.

This is aligned with the observation that overall change in binding free energy upon mutation is favorable. The sRBD is more stable upon binding and specifically the residues at the interface of ACE2 and RBD that provide increased stability (decreased flexibility) upon binding in the mutant. The residues distal to the binding interface are actually more flexible upon binding. Although ACE2 is less flexible in WT, RBD is more flexible. This corresponds to the NMA and DCCM data that the mutant system has more fluctuations at the distal ends of each protein with a stable hinge at the ACE2:RBD interface. The distal regions do not exhibit comparable fluctuations in WT and the whole system is more rigid due to the RBD to ACE2 correlations observed via DCCM as well as the hydrogen bonds at either end of the interface that latch it together.

## Conclusions

We have used all-atom MD simulations of sRBD WT and N501Y mutants, their complexes with sACE2, and sACE2 alone, to present a possible explanation for the observations that the N501Y mutation results in a less favorable enthalpy of binding, a more favorable entropy of binding, and an overall more favorable binding free energy, compared to wild-type sRBD. Our analysis began with MM/GBSA calculations confirming the enthalpic effect, which we then explained by exhaustive cataloging of inter-protein H-bonding. We argued that the general trend in the change of H-bonding is toward residue pairs that are more on the periphery of the interface for the mutant. Cross-correlation and normal mode analysis revealed that the mutant complex displays more quaternary flexibility than the WT complex, which we argue is a direct result of the more “hinge-like” quality of the peripheral H-bond network in the mutant compared to WT. Both this increased large-scale flexibility and the observation that sACE2 fluctuations increase upon binding to mutant but not for WT together provide a rationale for the more favorable entropy change upon binding for the mutant compared to the wild-type sRBD.

Our results suggest that one route toward increased protein-protein binding affinity is altering the layout of interprotein contacts in the epitope region so that large-scale vibrational modes that do not compromise binding are allowed to remain active. We speculate that this could lead to the development of binding antagonists that act to suppress this flexibility, making a protein-protein interface more fragile. Further elucidating this hypothesis could be done by some selected point mutations; for example, a K353M_sACE2_ mutation would disrupt the strong N501Y_sRBD_ to K353_sACE2_ bond in the mutant system. In addition the mutation of central residues on sACE2 such as E37K_sACE2_ and D30E_sACE2_ could enhance affinity by lengthening the binding interface. Additionally, it is interesting to note that some of the interactions identified in this work implicate residues that have emerged in the recent Omicron variants including Y505H, K417N, and G496S. ^17^ These residues are all central interactions in the WT data presented here but could present as weaker binders to their ACE2 partners in their mutant forms. Reversing these mutations systematically back to WT could further elucidate the effect of peripheral and central interactions with a spotlight on these contemporary mutations.

## Acknowledgments

Funding from the National Institutes of Health (Grant No. P01 AI150471) is gratefully acknowledged. This work used the Extreme Science and Engineering Discovery Environment (XSEDE), which is supported by National Science Foundation grant number ACI-1548562. This work also used the Picotte cluster in the Drexel University Research Computing Facility. M. Gatchel was supported on a Research Experience for Undergraduates Site Project (NSF Award No. 1949718, PI: M. Soroush).

